# TruePaiR: Software for the Accurate Identification of Complementary piRNA Read Pairs in High-Throughput Sequencing Data

**DOI:** 10.1101/167452

**Authors:** Patrick Schreiner, Peter W. Atkinson

**Affiliations:** Interdepartmental Graduate Program in Genetics, Genomics & Bioinformatics, University of California, Riverside, CA 92521, USA; Department of Entomology and Institute for Integrative Genome Biology, University of California, Riverside, CA 92521, USA

## Abstract

piRNAs and their biogenesis pathways are well-conserved in Metazoans (Grimson et al. 2008). piRNAs have been implicated in transcriptional, post-transcriptional, and translational regulation (Grivna et al. 2006; Lin & Yin 2008; Brennecke et al. 2008; Brennecke et al. 2007; Aravin et al. 2007). We analyze the signatures of a critical process in the primary and secondary mechanism of piRNA biogenesis, referred to as the amplification loop.

The presence of U-1 and A-10 bias within piRNA populations is an indicator, but not an absolute measure of piRNA amplification. By further considering imperfect and perfect sequence complementarity within the first ten base pairs of piRNAs, the active site promoting secondary piRNA biogenesis, we developed practical and statistically powerful metrics to observe relative piRNA amplification. TruePaiR is a fast and effective general software tool to assess the relative utilization of piRNA amplification in high throughput sRNA sequencing data.

The results of TruePaiR runs in seven species and five tissues serve as a benchmark for meaningful context of piRNA amplification. The TruePaiR metrics provide foundational data regarding the in terms of species specificity, tissue specificity, as well as the relative participation based upon origin-based piRNA subsets regarding piRNA amplification. The low degree of variability of same sample TruePaiR runs allows for metric reliability, reproducibility, as well as the ability to detect subtle differences in piRNA amplification within and between species and tissues. Given that TruePaiR serves as an effective and consistent metric of piRNA amplification across species, it can represent a new, meaningful standard in the degree of piRNA amplification in a specific organism and tissue that is or is not expected to undergo piRNA amplification.

## Introduction

piRNAs are the largest, in both size and number, distinct subclass of sRNAs (Zhang et al. 2014). Yet, piRNA biogenesis, targeting, and function are less well-understood relative to other sRNA pathways: siRNAs and miRNAs. piRNAs are quite distinct from the siRNA and miRNA pathways (Grimson et al. 2008; Aravin et al. 2007; Brennecke et al. 2007; Murchison & Hannon 2004).

piRNAs are noticeably distinct in that they are longer in sequence length, from 24-33 nts, relative to siRNAs and miRNAs (Aravin et al. 2007; Brennecke et al. 2007; Zhang et al. 2014). piRNAs are not known to form a hairpin secondary structure, and therefore have a Dicer-independent biogenesis (Aravin et al. 2007; Brennecke et al. 2007; Grimson et al. 2008). piRNA contain a 3’-O-methyl modification, modulated by HEN1, to protect from modification at the 3’ end of piRNAs (Horwich et al. 2007; Saito et al. 2007; Yang et al. 2006).

piRNAs are generated via a primary and secondary mechanism of biogenesis. Primary piRNAs derive from discrete genomic loci, referred to as piRNA clusters. piRNA clusters range from 5 to several hundred kbps in length and generally persist in heterochromatin (Arensburger et al. 2011; Brennecke et al. 2007). TE remnants are the major known component of piRNA clusters, which also can contain sequences of genic, viral, and unknown origin (Aravin et al. 2007; Schreiner & Atkinson 2017). Hundreds of millions of unique piRNA sequences have been identified, since piRNA sequences are not well-conserved amongst Metazoans (Zhang et al. 2014). piRNA cluster loci, however, are well-conserved by species (Schreiner & Atkinson 2017; Malone & Hannon 2010; Zanni et al. 2013; Malone & Hannon 2009; Grimson et al. 2008). piRNAs have been implicated in transcriptional, post-transcriptional, and translational regulation within the cell (Grivna et al. 2006; Lin & Yin 2008; Brennecke et al. 2008; Brennecke et al. 2007; Aravin et al. 2007).

Primary piRNA biogenesis is initiated via the transcription of a single, long piRNA precursor transcript (Brennecke et al. 2007). The Zucchini endonuclease slices the primary piRNA precursor molecule, generally resulting with a U at the first position of mature piRNAs (Nishimasu et al. 2012). Mature piRNAs then associate with the PIWI protein, Aub, to form a RNA-induced silencing complex (RISC) (Schwarz et al. 2004; Brennecke et al. 2007). The RISC is then guided to secondary piRNAs via complementarity of the associated primary piRNA.

Secondary piRNAs are generated as a result of the slicing mechanism of the RISC (Aravin et al. 2007). Argonaute, and therefore PIWI, proteins slice between the tenth and eleventh base pair of target molecules (Tolia & Joshua-Tor 2007). Initially, the Ping-Pong model of piRNA amplification suggested that given that adenine complements the uracil at the first position of the primary piRNA, and the secondary piRNA complements in the reverse orientation, the tenth position of secondary piRNAs generally have an A at position ten (Holbrook et al. 1991; Brennecke et al. 2007). An alternative model challenged this hypothesis, suggesting rather that the A-10 bias arises as a result of intrinsic preference of the target molecules of Aubergine (Wang et al. 2014). The 3’ end of the piRNAs trail on the opposite ends of the complex, and therefore, do not necessarily compliment (Zamore 2010; Aravin et al. 2007).

The degree of piRNA amplification is an important metric for assessing the activity of the piRNA biogenesis pathways. A metric exists to assess the degree of piRNA amplification in high throughput sRNA sequencing data considering the extent of the U-1 and A-10 bias. A Z-score test statistic can be calculated a to quantitate the significance of the observed bias within piRNA populations using the “pingpong” function to quantitate U-1 and A-10 bias overrepresentation within the NGS Toolbox of the piRNA cluster database (Zhang et al. 2011; Rosenkranz & Zischler 2012). Although, a method has not been developed to consider the sequence complementarity of piRNAs within a sRNA dataset of interest.

In order to correctly specifically identify sRNA pairs that have the potential to complement, the sequences of the piRNAs must be considered for compatibility. TruePaiR uses sequence complementarity to detect read pairs that are likely to facilitate Ping-Pong amplification in sRNA high throughput sequencing data.

## Materials and Methods

### Workflow

sRNA reads are the required input for TruePaiR in FASTA or FASTQ format. piRNAs are distinguished from other sRNAs using a length threshold greater than 23 nucleotides. Under the current model of Ping-Pong amplification loop, piRNA base pairs 11 and beyond don’t facilitate complementarity. Therefore, piRNA reads are trimmed to include only the first ten base pairs of the piRNAs. piRNA reads are then binned by those exhibiting only a U at the first position, those exhibiting only an A at the tenth position, and those exhibiting both a U at the first position and an A at the tenth position. Reads without a piRNA signature are not considered in assigning sRNA read pairs.

Mapping is then performed using Bowtie2 on the three partitions to predict complementary piRNA reads: (1) U-1 piRNAs as the subject and A-10 piRNAs as the reference set (2) U-1 piRNAs as the subject and piRNAs with both U-1 and A-10 as the reference set, and (3) A-10 piRNAs as the subject and piRNAs with both U-1 and A-10 as the reference set (Figure 1).

**Figure 4.1.**
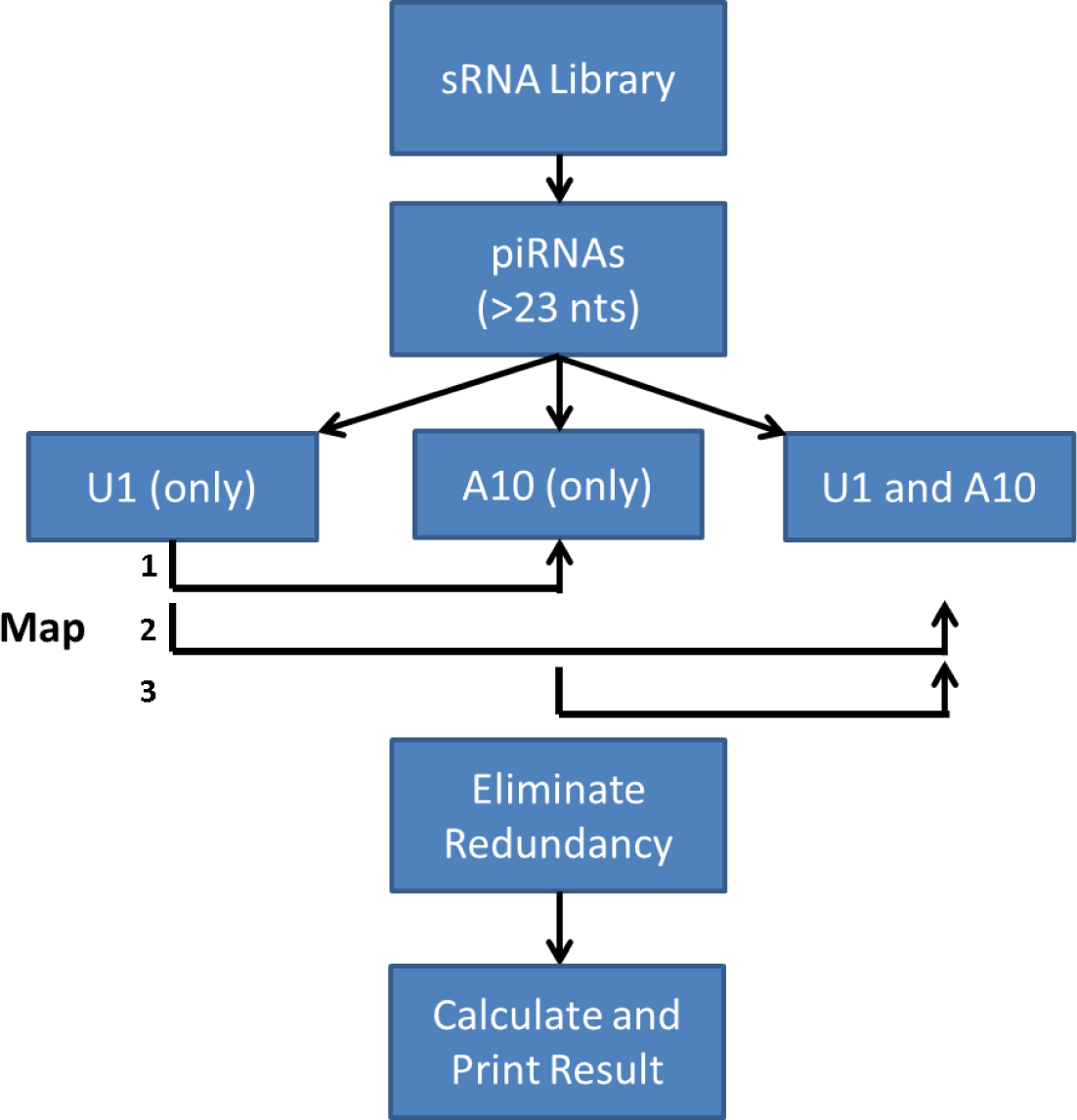
TruePaiR Workflow. A depiction of the steps that are utilized in the assessment of relative piRNA amplification using TruePaiR.

A verbose option is available to maintain intermediate files for downstream analysis. The intermediate files can give insight into the only U-1, only A-10, and both U-1 and A-10 subsets, as well as the mapping metadata associated with the TruePaiR run.

The three resulting SAM files, from each mapping run, are appended into a single file. Redundancy is removed within the file to be certain that each read is associated or not associated with a pair a maximum of one time. That is, each piRNA has a binary state in the TruePaiR algorithm: zero if the piRNA has no piRNA complement and one if the piRNA has at least one piRNA complement.

TruePaiR reports metrics regarding the number of piRNAs and the percentage of piRNAs with a U at the first position, an A at the tenth position, piRNAs that have a *possible* piRNA complement (0-2 mismatches), and piRNAs that have a *perfect* piRNA complement.

## Results

### Software Performance

TruePaiR is written and executed using R software. When the number of piRNA reads was under ten million, TruePaiR consistently completed under 10 minutes. However, the timing of TruePaiR runs can vary based upon library size and degree of complementarity of the piRNAs.

### Benchmarking Degree of piRNA Amplification

In order to appropriately interpret the TruePaiR output, we established benchmark values based on five tissues within seven species that have known or implicated activity in piRNA amplification (Brennecke et al. 2007; Aravin et al. 2007). Benchmarking using model organisms, and tissues with known piRNA pathway activity, serves to assess the ability of TruePaiR to make *a posteriori* assessments regarding the utilization of the Ping-Pong pathway using piRNA reads.

Variability of TruePaiR metrics was relatively low amongst same tissue samples within the same species. On average in the species and tissues observed in this analysis, the standard deviation relative to the observed values for U-1 presence in piRNAs was 9.5%, A-10 presence was 7.8%, possible pairs was 11.3%, and perfect pairs was 24.7% (Figure 4.2). The numbers of piRNAs in each library varied from 269,297 to 77,761,751 with an average of 9,498,527 and median of 4,674,392 piRNAs per library.

**Figure 4.2.**
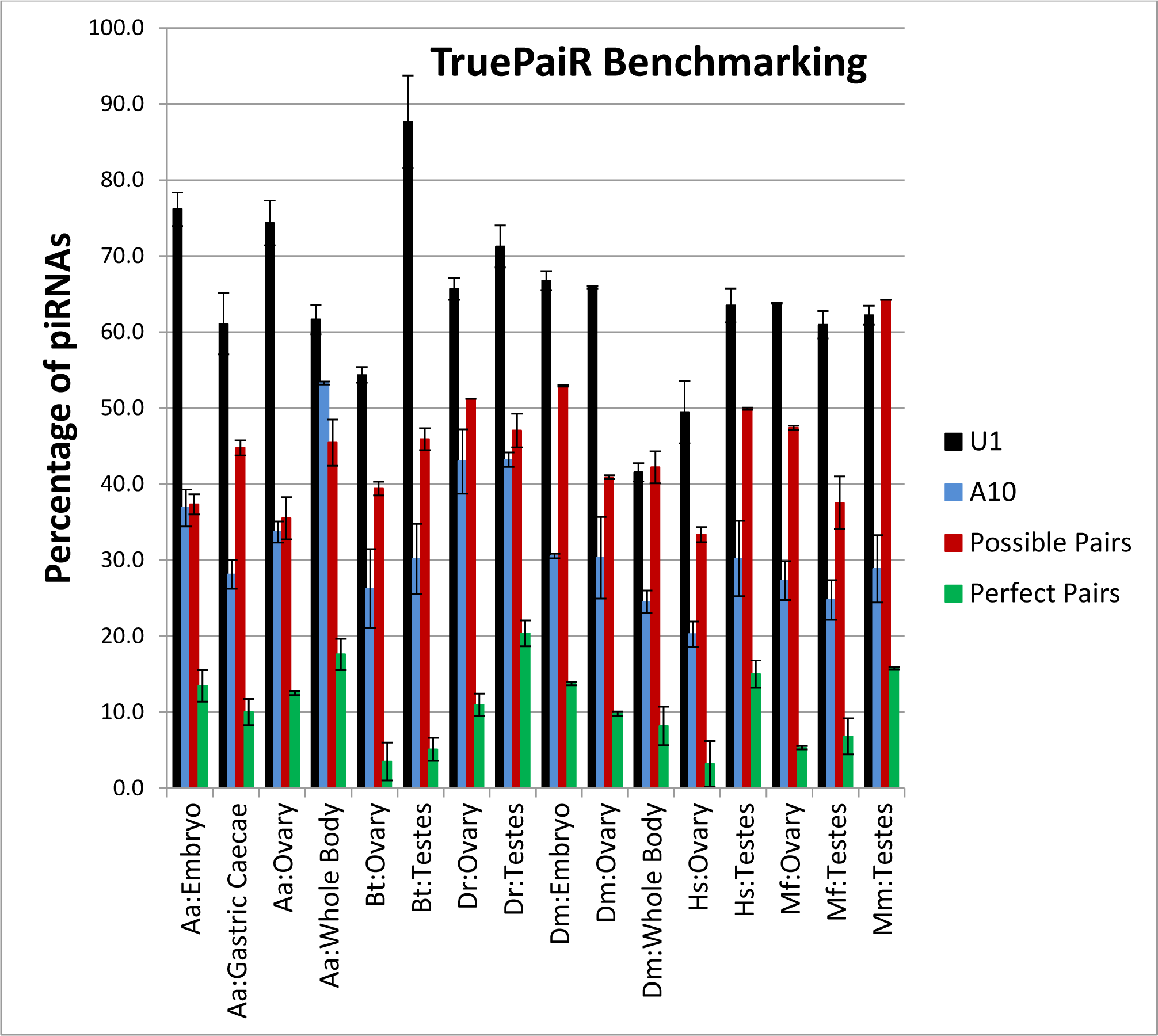
TruePaiR Benchmarking. Representation of the TruePaiR metrics oberseved in seven species and five tissues with replicate libraries. TruePaiR metrics include the percentage of piRNAs with a U at the first position (black), an A at the tenth position (blue), at least one possible piRNA complement (0-2 mismatches - red), and at least one perfect piRNA complement (green).

### Species-specific piRNA Amplification

TruePaiR metrics within the same tissue highlighted fundamental differences in piRNA populations and amplification among Metazoans. In ovarian tissue, the degree of U-1 bias within the piRNAs did not significantly differ between *Drosophila melanogaster, Danio rerio*, and *Macaca fascicularis*. Aedes aegypti had a significantly higher, while Bos Taurus and Homo sapiens had a significantly lower degree of U-1 bias in the piRNAs. The degree of A10 bias differed significantly between *Danio rerio, Aedes aegypti, Drosophila melanogaster, Macaca fascicularis, Bos taurus*, and *Homo sapiens* from greatest to least in A-10 representation amongst the piRNAs.

piRNA amplification, defined with indefinite sequence complementarity, was not significantly different between *Aedes aegypti, Bos taurus, Drosophila melanogaster*, and *Homo sapiens*. The potential for imperfect piRNA complementarity was significantly higher in *Danio rerio* and *Macaca fascicularis*. piRNA amplification with perfect sequence complementarity was significantly lower in each species relative to imperfect pairing. *Aedes aegypti, Danio rerio*, and *Drosophila melanogaster* had significantly more potential for perfect piRNA complements relative to *Bos taurus* and *Homo sapiens* (Figure 4.3A).

**Figure 4.3.**
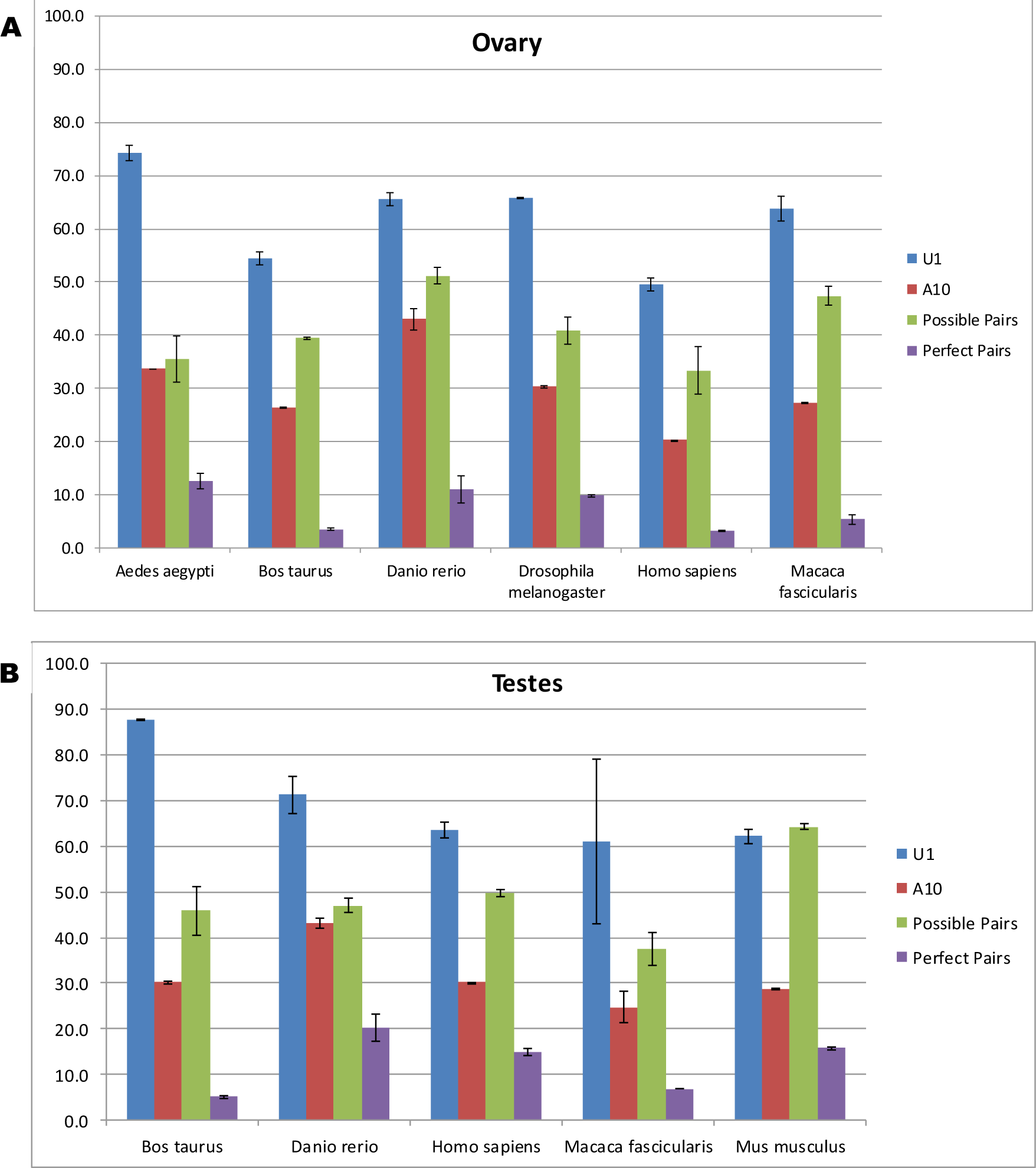
Species-specific Degree of piRNA Amplification. Histogram of the TruePaiR results for **(A)** ovary and **(B)** testes samples across species.

In testes tissue, the degree of U-1 bias did not significantly differ between *Homo sapiens, Macaca fascicularis*, and *Mus musculus*. piRNAs from *Bos taurus* testes were significantly higher in the degree of U-1 bias relative to the other species observed. The U-1 bias varied greatly (59.3%) *Macaca fascicularis* testes piRNAs. The degree of A-10 bias did not significantly differ between *Bos taurus, Homo sapiens, Macaca fascicularis*, and *Mus musculus* testes samples.

Possible piRNA amplification was not significantly different in *Bos taurus, Danio rerio*, and *Homo sapiens. Macaca fasciularis* had significantly less, while *Mus musculus* had significantly more potential for possible piRNA pairs. Perfect complementarity in facilitating piRNA amplification varied significantly in testes samples across species. *Danio rerio, Mus musculus, Homo sapiens, Macaca fascicularis*, and *Bos taurus* demonstrated the greatest to least potential for perfect piRNA pairs (Figure 4.3B).

### Tissue-specific piRNA Amplification

The observation of TruePaiR metrics regarding relative piRNA amplification within different tissues of the same species allows for an objective assessment of piRNA amplification between tissues.

In *Aedes aegypti*, consistent with previous research in germline and somatic piRNA amplification, ovary and embryo tissue demonstrated a significantly higher degree of U-1 bias in the piRNAs relative to gastric caecae and whole body (Brennecke et al. 2007; Aravin et al. 2007). The degree of A-10 piRNA bias differed significantly between whole body, embryo, ovary, and gastric caecae from greatest to least. The number of perfect and imperfect possible pairs did not significantly differ bwteen the tissue (Figure 4.4A).

**Figure 4.4.**
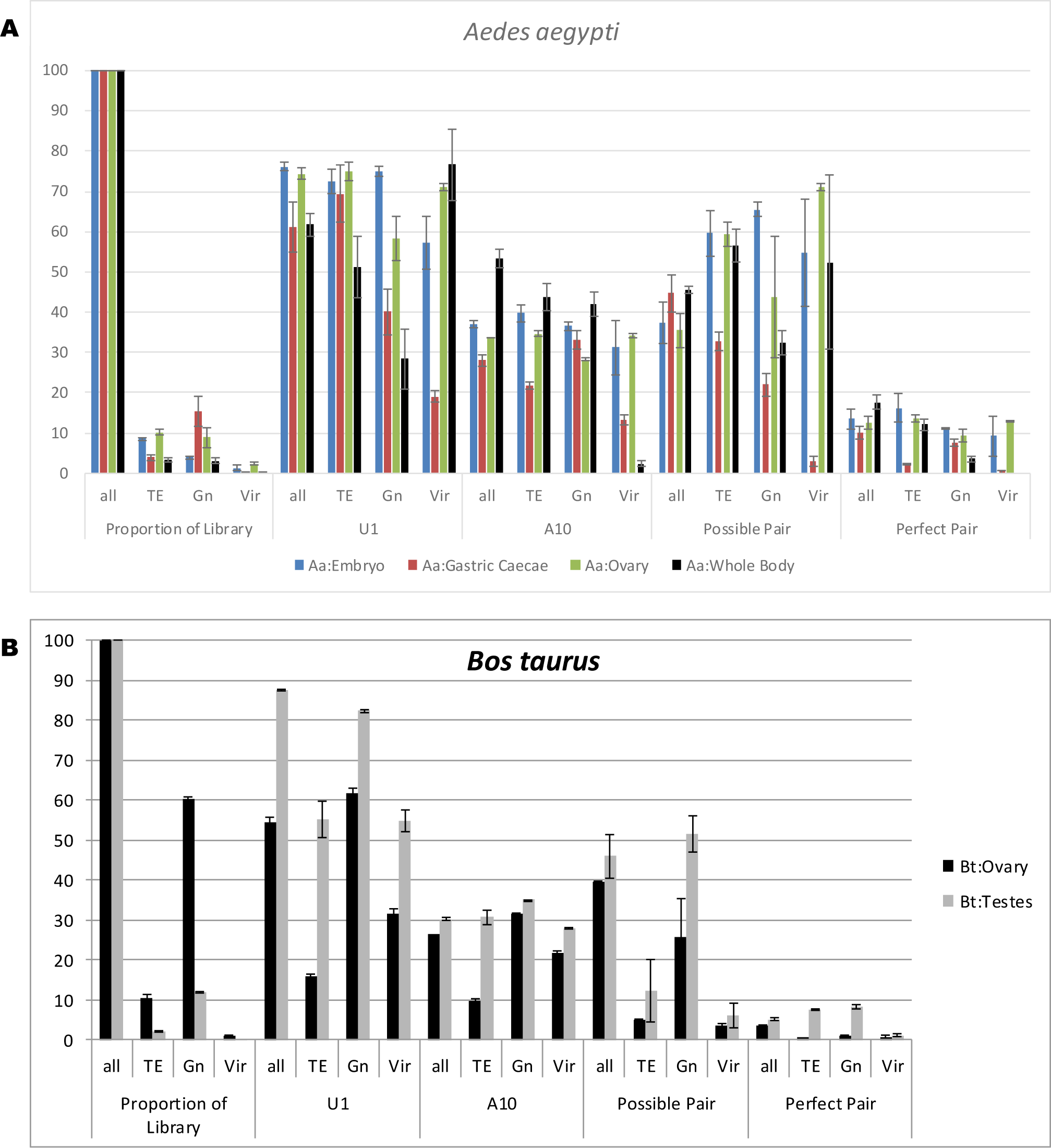
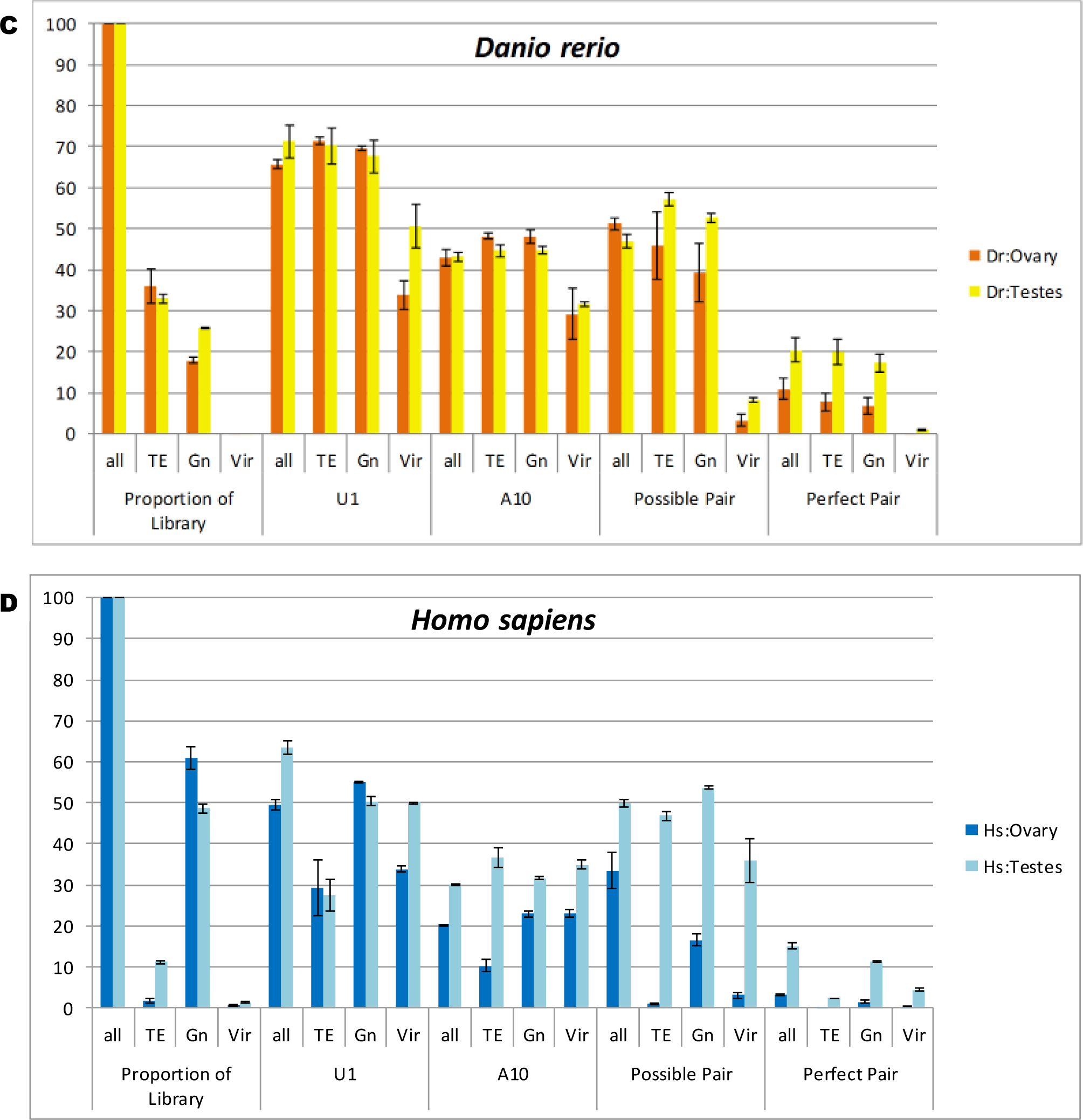
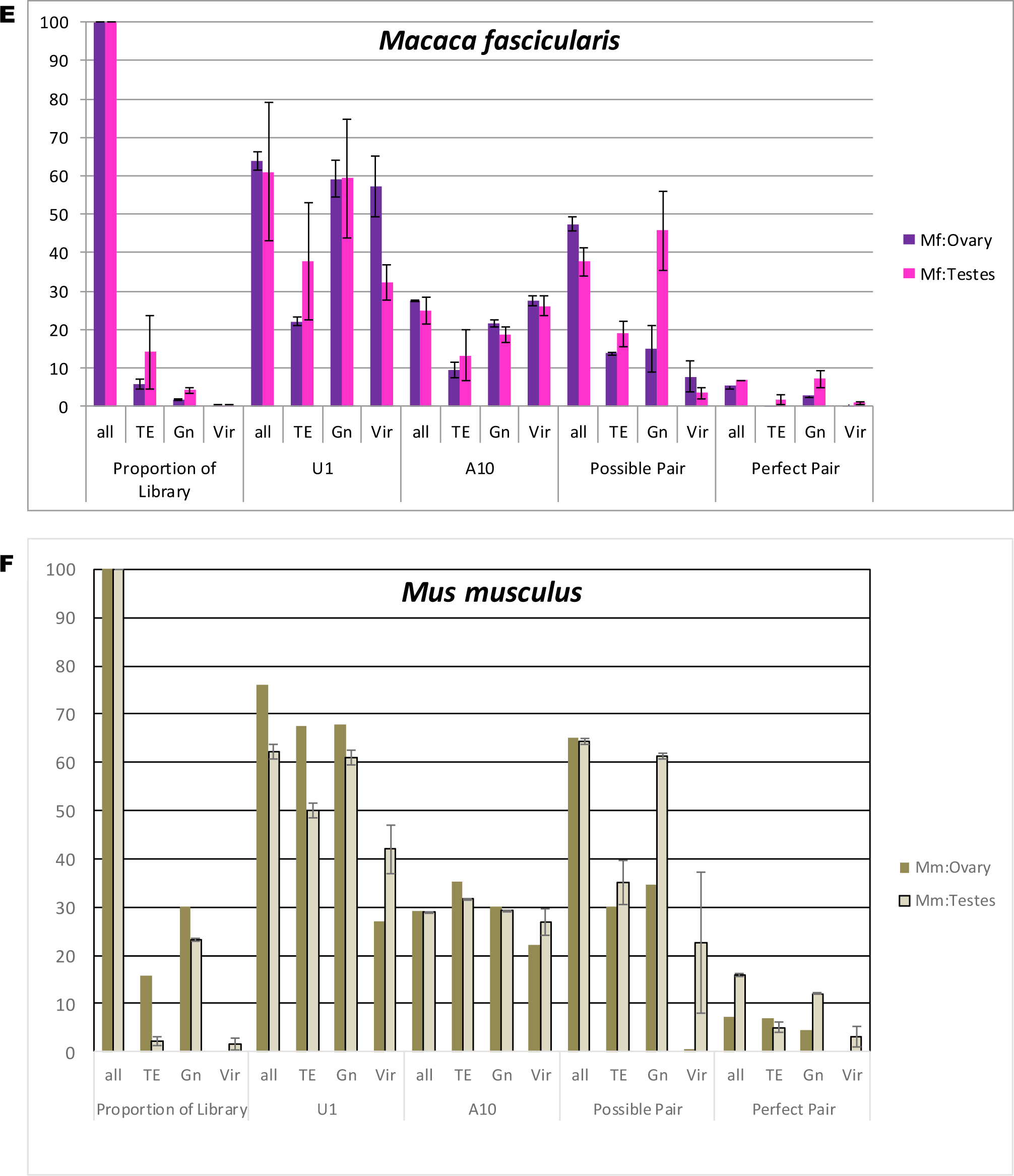
Tissue-specific Degree of piRNA Amplification. Histogram representing the differential proportions of piRNAs with U-1, A-10, a possible piRNA pair, and a perfect piRNA pair within ovary and testes tissues of **(A)** *Aedes aegypti* **(B)** *Bos taurus* **(C)** *Danio rerio* **(D)** *Homo sapiens* **(E)** *Macaca fascicularis* and **(F)** *Mus musculus*.

In *Bos taurus*, testes demonstrated a significantly higher degree of U-1 bias, A-10 bias, possibility of imperfect pairs, and possibility of perfect pairs relative to ovarian tissue (Figure 4.4B).

In *Danio rerio*, ovarian tissue exhibited a significantly greater degree of U-1 bias relative to testes. No significant difference was observed between the degree of A-10 bias between tissues. Ovarian tissue demonstrated a higher potential of imperfect piRNA pairs, but less of a potential for perfect pairs relative to testes (Figure 4.4C).

In *Homo sapiens*, testes tissue exhibited a significantly higher degree of U-1 bias, A-10 bias, possibility of imperfect pairs, and possibility of perfect pairs relative to ovarian samples (Figure 4.4D).

In *Macaca fascicularis*, no significant difference was observed in the degree of U-1 nor A-10 bias. Ovarian tissue demonstrated a significantly greater degree of possible piRNA complements, but a significantly lower degree of perfect piRNA complements (Figure 4.4E).

In *Mus musculus*, replicate libraries of piRNAs from ovarian tissue were not available to assess deviation between samples. However, piRNAs from ovarian tissue demonstrated a higher degree U-1 bias relative to testes samples. The difference in the degree of A-10 piRNA bias and imperfect pairing was consistent between tissues. Although, testes piRNAs exhibited a higher proportion of perfect piRNA pairs relative to ovarian tissue (Figure 4.4F).

### piRNA Origin and Relative Amplification

Further, observing subsets of the piRNAs in a particular library by their sequence of origin can provide insight into the nature of the piRNAs that are facilitating piRNA amplification. All piRNAs represent metrics gathered from sRNA reads greater than 23 nucleotides in length. TE-derived piRNAs were determined by piRNA homology to TEs available in the RepBase database (Jurka et al. 2005). Gene-derived piRNAs were determined by transcript reference datasets respective to the species under observation. Virus sequences were extracted from the NCBI database (Sayers et al. 2011).

Relative amplification based upon piRNA origin varied greatly between species and tissues. However, TE- and gene-derived piRNAs were consistently a more prevalent subset of the total piRNA population and had a higher capability of participation in piRNA amplification in the species and tissues observed relative to viral-derived piRNAs (Figure 4.4).

### Application in piRNA Pathway Knockdowns (KDs)

Data available within the Short Read Archive and Gene Expression Omnibus is available for heterozygous and knockdown conditions of several proteins that are critical in promoting piRNA amplification (Leinonen et al. 2010; Edgar et al. 2002). Most notably, we observed knockdown effect on the piRNA populations of Piwi, Aubergine, Zucchini, and Argonaute 3.

Although replicate libraries were not available to establish statistical significance, a distinction can be noted between heterozygous and knockdown libraries in *Drosophila melanogaster* ovaries (Malone et al. 2009). In Piwi KD, the proportion of piRNA populations with a U-1 bias was higher by 45.4%, A-10 bias was lower by 18.4%, possible pairs was higher by 44.2%, and perfect pairs was higher by 3.0% in control libraries relative to KD. In Aubergine KD, the proportion of piRNA populations with a U-1 bias increased by 41.3%, A-10 bias decreased by 23.0%, possible piRNA pairs increased by 11.9%, and perfect piRNA pairs increased by 6.6%. In Zucchini KD, the proportion of piRNA populations with a U-1 bias increased by 6.5%, A-10 bias increased by 1.9%, possible piRNA pairs increased by 3.4%, and perfect piRNA pairs increased by 5.1% (Figure 4.5).

**Figure 4.5.**
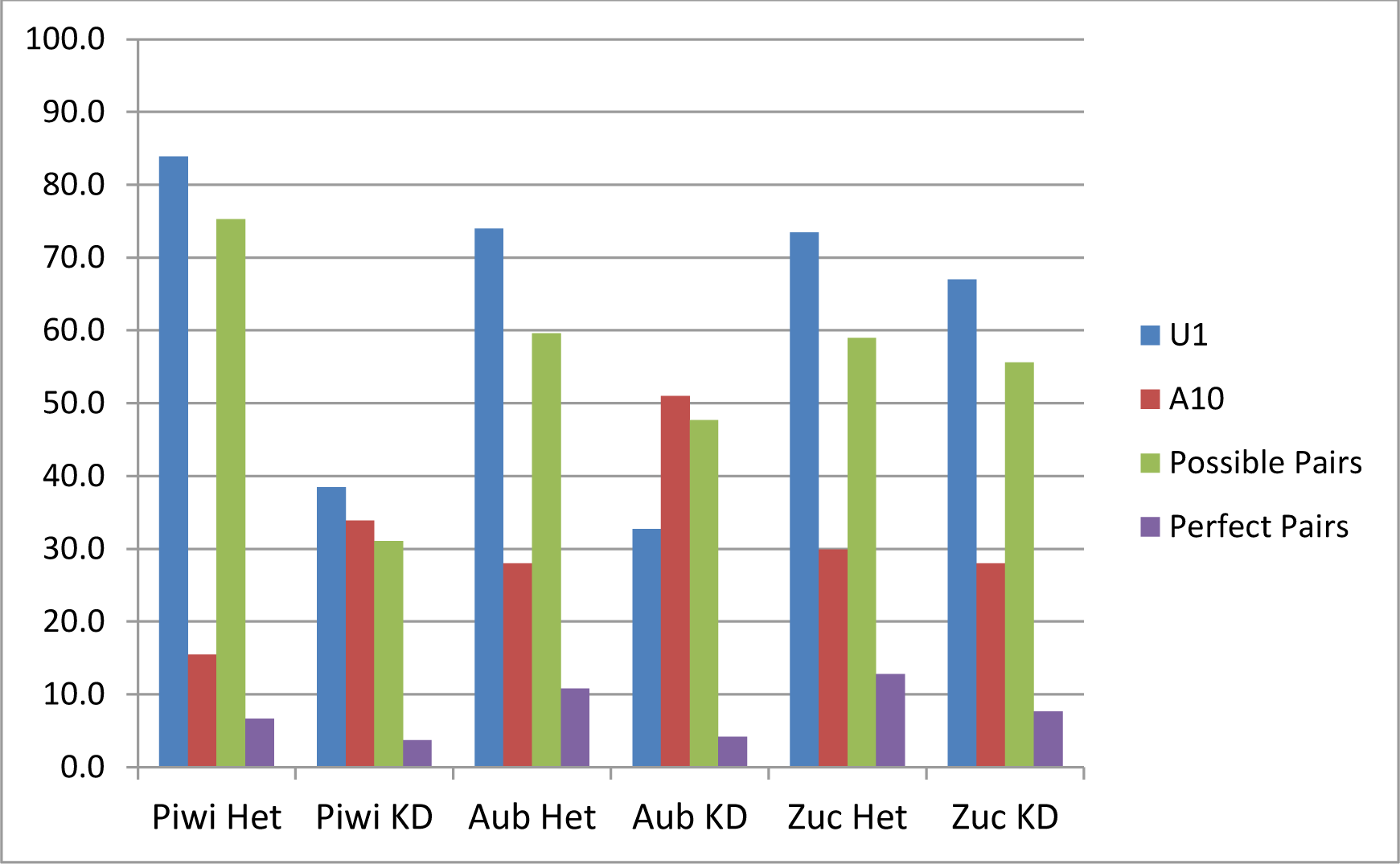
Relative piRNA Amplification in Available KD Libraries. Observation of TruePaiR metrics in publicly available Piwi, Aubergine (Aub), and Zucchini (Zuc) heterozygous and knockdown ovary in *Drosophila melanogaster* (Malone et al. 2009).

Further, piRNAs from male and female fourth instar larvae in *Aedes aegypti* in the presence and absence of AGO3 (Han and Atkinson, unpublished). The model of piRNA biogenesis suggests that AGO3 is a critical protein involved in the promotion of piRNA amplification (Brennecke et al. 2007). Upon successful AGO3 KD in males, a reproducible and statistically significant difference is observed in the degree of piRNA amplification relative to uninduced male fourth instar larvae. However, due to an explanation that is still under investigation, AGO3 transcript was not suppressed in female fourth instar larvae upon induction (Han and Atkinson, unpublished). TruePaiR detected no significant difference of piRNA amplification in female fourth instar larvae (Figure 4.6).

**Figure 4.6.**
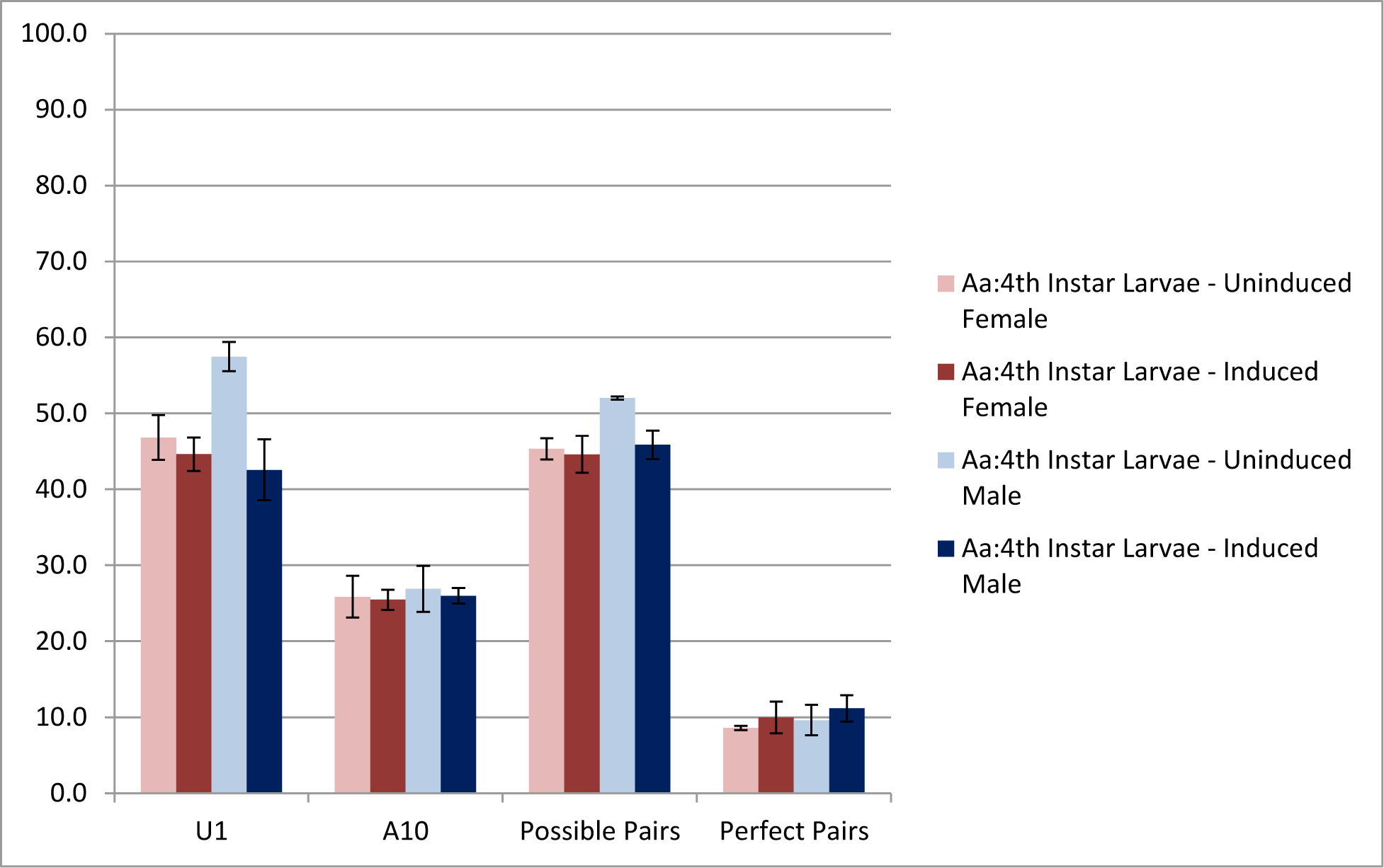
Relative piRNA Amplification in Ago3 KD. TruePaiR runs in *Aedes aegypti* fourth instar larvae males and females in the presence (uninduced) and absence (induced) of Ago3.

## Discussion

The presence of U-1 and A-10 bias within piRNA populations is an indicator, but not an absolute measure of piRNA amplification. By further considering imperfect and perfect sequence complementarity within the first ten base pairs of piRNAs, the active site promoting secondary piRNA biogenesis, we developed practical and statistically powerful metrics to observe relative piRNA amplification (Brennecke et al. 2007).

TruePaiR is a fast and effective tool to determine the relative utilization of the piRNA amplification using high-throughput sRNA sequencing data. TruePaiR is an effective and robust gauge of the piRNA amplification using a single sRNA subpopulation.

The TruePaiR results were accurate in detecting relatively high piRNA amplification in tissue, such as ovary and testes, that has been well-studied to participate in the piRNA pathway. Further, TruePaiR was able to detect piRNA amplification in *Aedes aegypti* gastric caecae and fourth instar larvae, which are tissues that have not been previously known to utilize this pathway (Han and Atkinson, unpublished). TruePaiR was also capable of detecting low levels of piRNA amplification in species, tissues, and piRNA subsets in which the secondary pathway of piRNA biogenesis has little or no activity.

Established benchmark values, in model species and tissues known to undergo piRNA amplification, allow for the observation of meaningful context of the TruePaiR metrics for species or tissue in which the degree of piRNA amplification is not well-understood (Figure 4.2-4.3). The results presented herein provide foundational data regarding piRNA amplification in terms of species specificity, tissue specificity, as well as the relative participation based upon piRNA origin. General trends in the proportion of U-1 piRNAs, A-10 piRNAs, and number of piRNA complements are consistent with conserved model of piRNA biogenesis via the amplification loop (Brennecke et al. 2007). The TruePaiR benchmark values characterize the difference in relative piRNA amplification, which can lead to downstream experimentation to identify species- or tissue-specific factors that affect piRNA biogenesis.

Sample variation was minor in independent sRNA samples of the same tissue. The detected differences in the TruePaiR metrics between species and tissues may be due to species-specific factors that facilitate or inhibit piRNA amplification, the number of active piRNA clusters, the number of generated piRNAs, or the sequence content of generated piRNAs (Figure 4.2-4.3). Even considering the innate variability between organisms and library preparations, the results of TruePaiR were very consistent in assessing piRNA amplification in particular species and tissues. Consistency across same sample TruePaiR runs allows for a reliable and reproducible assessment of relative piRNA amplification.

TruePaiR demonstrated capability to detect differences in relative piRNA amplification between conditions. Differences were observed between hetereozygous and knockdown sRNA libraries of Piwi, Aub, and Zucchini, in a similar magnitude as previously described, while providing specific metrics regarding the effects of each particular knockdown (Malone et al. 2009) (Figure 4.5). Further, the TruePaiR metrics of possible piRNA pairs was capable of distinguishing, with both reproducibly and statistical significance, minor differences in piRNA amplification in triplicate sRNA libraries of *Aedes aegypti* fourth instar larvae upon Ago3 control and knockdown (Han and Atkinson, unpublished) (Figure 4.6).

The TruePaiR results showed consistently low levels of perfect piRNA pairs within the first ten base pairs, even in tissue that are known to have the highest levels of piRNA amplification. The relative proportion of possible piRNA pairs, allowing up to two mismatches in the first ten base pairs, increased significantly in germline tissue known to be involved in piRNA amplification. (Figure 4.4). These results support a piRNA amplification model of imperfect complementarity in the first ten base pairs of piRNA complements.

Given that TruePaiR serves as an effective and consistent metric of piRNA amplification across species, it can represent a new, meaningful standard in the degree of piRNA amplification in a specific organism and tissue that is or is not expected to undergo piRNA amplification.

